# Confident Identification and Quantification of Mouse Brain Tissues Reveals Sirtuin 5-Dependent Regulation

**DOI:** 10.64898/2026.05.26.726073

**Authors:** Landgrave-Gomez Jorge, Bons Joanna, Vega-Hormazabal Genesis, Riley Rebeccah, Schilling Birgit, Verdin Eric

**Affiliations:** Buck Institute for Research on Aging, Novato, California 94945, USA

**Keywords:** methylmalonylation, proteomics, data-independent acquisition (DIA-MS), lysine acylation, SIRT5, myelin basic protein

## Abstract

Methylmalonylation is a non-enzymatic lysine post-translational modification derived from methylmalonyl-CoA, a reactive intermediate that accumulates during mitochondrial dysfunction and branched-chain amino acid catabolism. Although reported in models of methylmalonic acidemia, its broader distribution and functional relevance remain largely unexplored. Progress has been hindered by a key analytical challenge: methylmalonyl-and succinyl-lysine are isobaric (+100.0160 Da) and generate overlapping mass spectrometric fragmentation spectra, preventing confident identification in conventional proteomic workflows.

Here, we establish a straightforward proteomic workflow that overcomes this barrier and enables confident identification and quantification of lysine methylmalonylation by combining antibody-based enrichment with data-independent acquisition mass spectrometry (DIA-MS). Anti-malonyl antibodies were used to enrich methylmalonylated peptides through cross-reactivity. Using synthetic peptide standards containing malonyl-, succinyl-, or methylmalonyl-lysine, we defined distinguishing analytical features including chromatographic retention time, ion mobility, and fragmentation patterns.

Applying this approach to mouse brain tissues from Sirtuin-5 (SIRT5) knockout and wild-type mice, we identified 44 methylmalonylated peptides across 41 proteins, enriched in neuronal and myelin-associated proteins (NEFM, NEFL, MBP) and mitochondrial enzymes such as ADT1. Several sites were increased in SIRT5-deficient brains, consistent with regulation by this mitochondrial deacylase. Functional assays demonstrated that methylmalonylation of myelin basic protein (MBP) impairs lipid binding, linking this modification to myelin stability.

Together, this workflow enables confident methylmalonylation identification and defines it as a widespread and regulated modification in the brain, providing a framework to study metabolically driven protein acylation in neurobiology and disease.

**Significance:** Lysine methylmalonylation has remained largely unexplored due to its isobaric overlap with succinylation, which prevents confident identification using conventional proteomic workflows. Here, we establish an integrated strategy combining antibody-based enrichment, data-independent acquisition mass spectrometry, and orthogonal analytical features to resolve these modifications with high confidence. Applying this approach to mouse brain tissue reveals a SIRT5-regulated methylmalonylome enriched in mitochondrial and myelin-associated proteins, including myelin basic protein (MBP). Functional assays demonstrate that methylmalonylation impairs MBP lipid binding, linking this modification to myelin stability. Beyond this specific application, our workflow provides a generalizable framework to resolve isobaric post-translational modifications and expands the study of metabolically driven protein acylation in neurobiology and disease.

## Introduction

Lysine acylations are a diverse class of post-translational modifications (PTMs) that modulate protein function by the reversible covalent addition of acyl groups derived from cellular metabolism ^1,2^. Among these, negatively charged lysine acylations—including malonylation, succinylation, and the recently described methylmalonylation—are tightly linked to mitochondrial dysfunction and metabolic stress ^3–6^. They arise from reactive short-chain acyl-CoA species and accumulate when CoA metabolism or NAD⁺ homeostasis is impaired, as occurs in inborn errors of metabolism (IEMs) and during aging ^7–10^. Owing to their negative charges, these lysine acylations shift lysine residue charge from +1 to -1, which can substantially alter protein electrostatics and disrupt protein–protein and protein–lipid interactions ^11–13^.

Methylmalonylation originates from methylmalonyl-CoA, a byproduct of branched-chain amino acid (BCAA) and odd-chain lipid catabolism ^9,14,15^ (Fig. 1A). These metabolic pathways are particularly active in the brain, where oligodendrocytes rely on lipid synthesis to sustain myelin production and excitatory neurons use BCAA catabolism to generate glutamate ^16–18^. Despite this vulnerability, the occurrence and function of methylmalonylation in the central nervous system remain poorly characterized. While methylmalonyl-lysine has been reported in liver tissue from methylmalonic acidemia models, broader tissue mapping has been hindered by analytical limitations ^6^.

**Figure 1.**
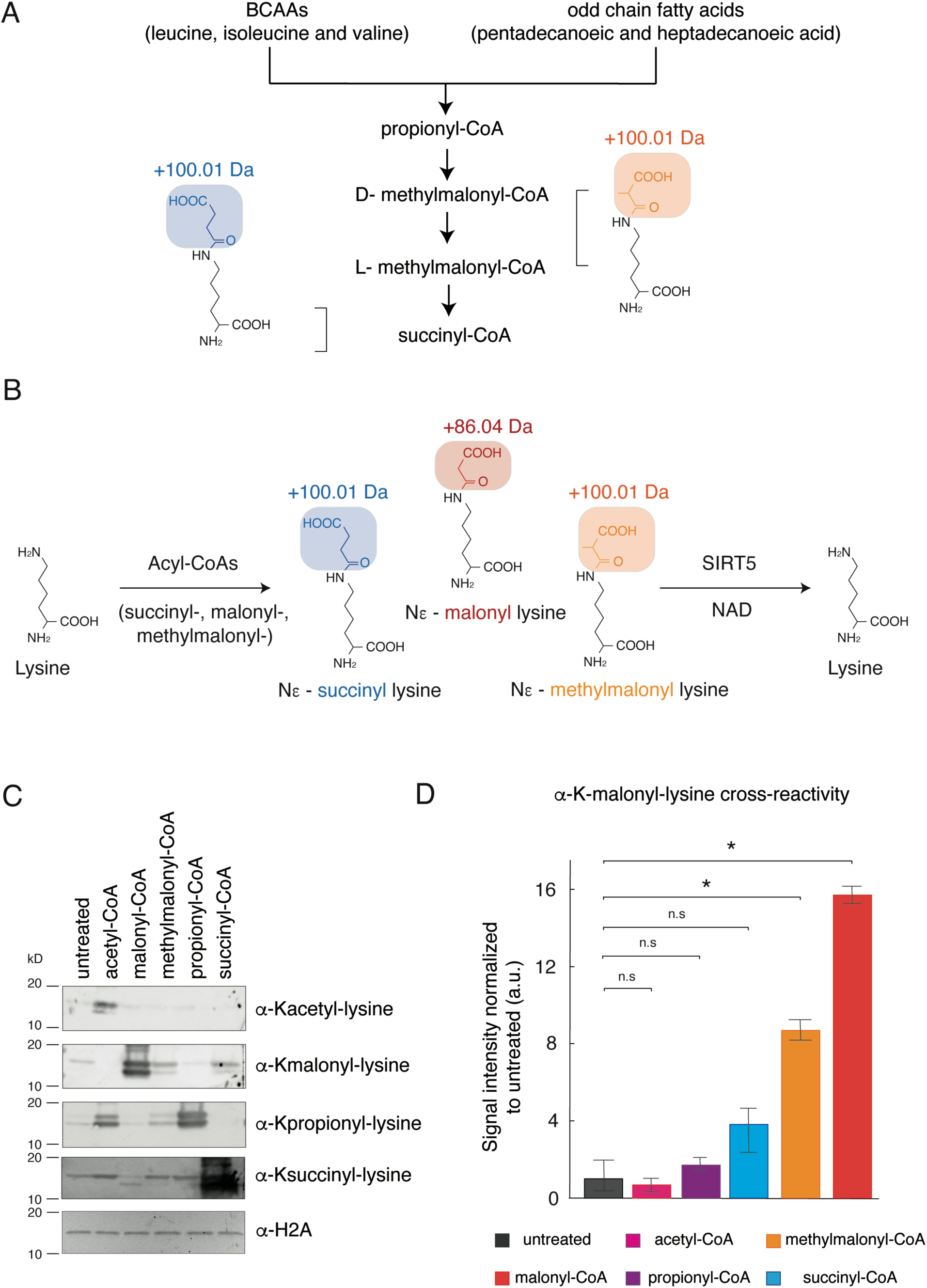
Chemical similarity and antibody cross-reactivity between lysine acylations. (A) Metabolic pathways generating acyl-CoA donors for lysine acylations. Branched-chain amino acid (BCAA) catabolism and odd-chain fatty acid oxidation produce propionyl-CoA, which is converted to methylmalonyl-CoA and subsequently to succinyl-CoA. These metabolites serve as donors for lysine acylations including methylmalonylation (in orange) and succinylation (in blue). (B) Lysine acylation occurs by the addition of an acyl group to the ε-amine of their side chain. This modification is reversible and can be removed by the NAD^+^-dependent deacylase Sirtuin 5 (SIRT5). The chemical structures and mass shifts of lysine acyl modifications examined in this study, including malonylation (+86.04 Da, in red), succinylation (+100.01 Da, in blue), and methylmalonylation (+100.01 Da, in orange), are represented. Succinyl- and methylmalonyl-lysine are isobaric, creating challenges for confident identification by mass spectrometry. (C) Immunoblot analysis demonstrating antibody cross-reactivity across different lysine acyl modifications. Recombinant histones were incubated with various acyl-CoA donors (acetyl-CoA, malonyl-CoA, methylmalonyl-CoA, propionyl-CoA, and succinyl-CoA) and probed with antibodies recognizing acetyl-, malonyl-, propionyl-, or succinyl-lysine. Ponceau stainings and H2A blots were used as loading controls. (D) Quantification of antibody signals (anti-acetyl-, malonyl-, propionyl-, or succinyl-lysine) in all acyl-CoA treatment conditions. Levels of acylation were normalized to the level of unmodified H2A. Mean and SD are represented (* p-value < 0.05).

Mass spectrometry-based proteomics has become a powerful tool for PTM analysis. Significant advances in modified peptide enrichment strategies, instrumentation capabilities, and computational pipelines have enabled comprehensive profiling and quantification of PTMs. However, analytical and bioinformatics challenges remain, among which the analysis of low abundance PTM for which no enrichment strategies are available, and the differentiation of isobaric/isomeric modified peptides. Both lysine methylmalonylation and lysine succinylation lead to a mass increase of +100.0160 Da (Fig. 1B), and these isobaric residues yield nearly indistinguishable fragmentation spectra, often resulting in misassignment or exclusion of these modifications in standard workflows ^5,6,19^. Moreover, no enrichment strategy is available that specifically target methylmalonylated peptides for downstream proteomic analysis. As a result, lysine methylmalonylation has likely been overlooked in existing acylome datasets, leaving its prevalence and biological relevance largely unexplored. Interestingly, anti-malonyl antibodies have been shown to cross-react with methylmalonylated epitopes, providing a route for enrichment and alleviate the problem of isobaric PTMs ^6,20^.

Here, we establish an integrated label-free proteomic workflow to address this gap, that combines anti-malonyllysine antibody-based methylmalonylated peptide enrichment with comprehensive data-independent acquisition-mass spectrometry (DIA-MS) for PTM identification, quantification, and site localization. We first validated the cross-reactivity of anti-malonyl lysine antibodies toward methylmalonyllysine epitopes using recombinant histones as a model, then used synthetic reference peptides with malonyl-, succinyl-, and methylmalonyl-lysine to characterize these PTMs and identify discriminating analytical features including peptide retention time in liquid chromatography, ion mobility separation, and diagnostic MS/MS fragmentation patterns. Applying this workflow to mouse brain tissues collected from wild-type and Sirtuin 5 (SIRT5) knock-out mice, a NAD^+^-dependent lysine de-malonylase and de-succinylase, we generated the first proteome-wide map of lysine methylmalonylation in the brain and identified myelin basic protein (MBP) as a prominent substrate whose methylmalonylation disrupted lipid-binding capacity in independent *in vitro* assays.

## Experimental Procedures

### Proteomic Sirt5-KO and WT mouse brain acylomes

We previously reported the quantitative analysis of protein malonylation and succinylation in Sirt5-KO and WT mouse brains (n = 4 each) ^19^. Briefly, whole brain tissues from 18-month-old mice were homogenized, trypsinized, desalted, and further enriched for malonylated or succinylated peptides using the PTMScan Malonyl-Lysine Motif Kit or PTMScan Succinyl-Lysine Motif Kit (Cell Signaling Technologies), respectively. Samples were analyzed in data-independent acquisition (DIA) mode on an Orbitrap Eclipse Tribrid platform (Thermo Fisher Scientific). DIA-MS data were analyzed using the library-free tool directDIA embedded in Spectronaut (version 14) and Skyline for acylated peptide identification and quantification and PTM site localization ^22^. Datasets can be retrieved using the MassIVE ID MSV000089606 and ProteomeXchange ID: PXD034275.

To investigate lysine methylmalonylation in brain tissues from Sirt5-KO and WT mice, we re-analyzed the DIA-MS data collected on malonylated peptide enrichments using directDIA (Spectronaut version 17.1.221229.55965). Data were searched against the same *Mus musculus* proteome (58,430 entries from UniProtKB-TrEMBL, accessed on January 31, 2018). Trypsin/P was set as digestion enzyme and four missed cleavages were allowed. Cysteine carbamidomethylation was set as fixed modification, and lysine methylmalonylation, methionine oxidation, and protein N-terminus acetylation as variable modifications. Data extraction parameters were set as dynamic. Identification was performed using 1% precursor and protein q-value. PTM localization was selected with and a localization probability cutoff of 0.75 was used. Quantification was based on the XICs of 3–6 MS2 fragment ions, specifically b- and y-ions, without normalization, as well as data filtering with q-value. Grouping and quantitation of PTM peptides were accomplished using the following criteria: minor grouping by modified sequence and minor group quantity by mean precursor quantity. Differential expression analysis was performed using a paired t-test, and p-values were corrected for multiple testing using the Storey method ^22^. Methylmalonylated peptides with p-value < 0.05 and absolute log_2_(fold-change) > 0.58 were considered significantly altered (Supplementary Table S1). The extracted fragment ion chromatograms (XICs) of the malonylated peptide LAADVG^147^**Kmalonyl**GSSQR, succinylated peptide LAADVG^147^**Ksuccinyl**GSSQR, and methylmalonylated peptide LAADVG^147^**Kmethylmalonyl**GSSQR from the mouse ADP/ATP Translocase 1 (UniProt ID: P48962) were visualized in Skyline ^21^, and their identification was supported by orthogonal validation using retention time, ion mobility, and fragmentation patterns from synthetic standards.

### Synthetic peptide standards

Synthetic malonylated, succinylated, and methylmalonylated peptides LAADVG^147^**Kacyl**GSSQR* from the mouse ADP/ATP Translocase 1 (UniProt ID: P48962) with a modified lysine at position K-147 were from Biosynth (>98% purity). Each modified peptide was synthetized as light form and heavy form containing stable isotope-labelled arginine residue (R* with ^13^C_6_, ^15^N_4_). Synthetic peptides were diluted in 2% acetonitrile (ACN), 0.1% formic acid (FA) in water for MS analysis.

### prm-PASEF acquisition and analysis

Synthetic peptides (10 pg each, corresponding to 7.7-7.8 fmol) were loaded onto Evotips Pure (Evosep) following the manufacturer’s protocol. Targeted assays were conducted on an Evosep One liquid chromatography system coupled to a timsTOF HT mass spectrometer (Bruker Daltonics). The solvent system consisted of 0.1% FA in water (solvent A) and 0.1% FA in ACN (solvent B). Peptides were eluted on a PepSep C_18_ analytical column (150 µm x 15 cm, 1.5 µm particle size; Bruker) using the 30 SPD method (44-min gradient length, 500-nL/min flow rate). A zero-dead volume emitter was installed in the nano-electrospray source (CaptiveSpray source, Bruker Daltonics), and the source parameters were set as follows: capillary voltage 1600 V, dry gas 3 L/min, and dry temperature 180°C. Each sample was acquired in parallel reaction monitoring coupled to parallel accumulation serial fragmentation (prm-PASEF) mode ^23^. TIMS accumulation was set to 150 ms and the ion mobility separation to 150 ms. The ion mobility ranged from 1/K_0_ = 0.60 to 1.60 Vs/cm^2^ and the covered m/z range was 100 to 1700. For the scheduled acquisition, the retention time range was 8.0-11.0 min and the mobility range (1/K_0_) 0.80-1.10 Vs/cm^2^, and the isolation window width was set to 3.0 Da. The collision energy was defined as a linear function of mobility, starting from 20 eV at 1/K_0_ = 0.6 Vs/cm^2^ to 59 eV at 1/K_0_ = 1.6 Vs/cm^2^. The prm-PASEF data were visualized with DataAnalysis (Bruker Daltonics) and processed with Skyline by extracting fragment ion chromatograms (XICs) ^22^.

#### GO enrichment

Proteins harboring high confidence methylmalonylation or succinylation sites were submitted to over-representation analysis against the *Mus musculus* background using STRING (version 12.0). Gene Ontology (GO) terms with q<0.05 were considered significant. Visualization used STRING (version 12.0).

#### Recombinant proteins and *in vitro* acylation

Histones (recombinant H2A/H2B/H3/H4; Epicypher; 15-0301, 15-0302, 15-0303 and 15-0304) and MBP (Active motif, 82617) were incubated with acyl-CoA donors (acetyl-, malonyl-, succinyl-, or methylmalonyl-CoA; Sigma-Aldrich; 0.2–1 mM) in 50 mM HEPES pH 8, 150 mM NaCl, 1 mM DTT, 2 mM MgCl₂ at 37 °C for 1–2 h. Reactions were desalted (Zeba spin columns) prior to downstream assays. Mean ± SD from three independent replicates is shown.

#### Immunoblotting

Proteins were resolved by SDS-PAGE and transferred to nitrocellulose. Membranes were blocked (5% BSA/TBST) and probed with anti-malonyl-lysine (PTM BIO; PTM-901, 1:1000), anti-propionyl-lysine (PTM BIO; PTM-201, 1:1000), anti-succinyl-lysine (PTM BIO; PTM-401, 1:1000), anti-acetyl-lysine (PTM BIO; PTM-132), anti-H2A (Cell Signaling; 2578, 1:1000 and anti-MBP (Cell Signaling; 78896, 1:1000) as indicated. Fluorescent secondary antibodies (Azure biosystems; SKUAC2128, SKUAC2129, 1:10000) were used. Band densities were quantified in ImageJ/Fiji and normalized to loading controls or Ponceau staining. Mean ± SD from three independent replicates is shown.

#### Lipid overlay (dot-blot) assays

PVDF membranes pre-spotted with myelin-enriched lipids (e.g., phosphatidylserine, phosphatidylinositol, sulfatide; Avanti 141101) were blocked (3% fatty-acid free BSA in TBS). MBP (unmodified or acylated) was incubated at 1 µg/mL for 60 min at RT. Bound MBP was fixed using 1% PFA and detected with anti-MBP (Cell Signaling; 78896, 1:1000) then quantified by densitometry. Signals were normalized to the unmodified/acyl control. Mean ± SD from three independent replicates is shown.

#### Data availability

The prm-PASEF data and methylmalonylated proteomic dataset have been uploaded to the Mass Spectrometry Interactive Virtual Environment (MassIVE) repository, developed by the Center for Computational Mass Spectrometry at the University of California, San Diego, and can be downloaded using the following FTP link: ftp://MSV000101474@massive-ftp.ucsd.edu or via the MassIVE website: https://massive.ucsd.edu/ProteoSAFe/dataset.jsp?task=5b263dff4b694de9bb848c4996f 9cc5a (MassIVE ID: MSV000101474; ProteomeXchange ID: PXD077235).

## Results

### Anti-malonyl antibodies enable enrichment of methylmalonylated peptides

A central challenge in studying protein methylmalonylation is the absence of antibodies specific for methylmalonyl-lysine, which has limited enrichment-based proteomics. Previous reports suggested that anti-malonyl antibodies may cross-react with methylmalonylated epitopes. To test this under controlled conditions, we used recombinant histones as a model substrate because of their high lysine density and susceptibility to non-enzymatic acylation. Histones incubated with malonyl-, succinyl-, or methylmalonyl-CoA were probed with anti-malonyl-lysine antibodies. As expected, malonylated histones yielded strong signals. Importantly, methylmalonylated histones were also robustly detected, whereas succinylated histones showed non-significant reactivity (Fig. 1C-D). These findings establish that anti-malonyl antibodies can recover methylmalonylated peptides while discriminating against succinylated epitopes.

### Synthetic peptide standards enable development of a workflow to discriminate methylmalonyl-from succinyl-lysine

Another challenge for confident methylmalonylation identification by mass spectrometry is the isobaric nature of methylmalonyl- and succinyl-lysine (+100.016 Da), leading to the generation of overlapping, multiplexed MS/MS fragmentation spectra. To further characterize these modifications, we synthesized three forms of the tryptic peptide LAADVG^147^**Kacyl**GSSQR* from the mouse ADP/ATP Translocase 1 (ADT1), each with a modified malonyl-, succinyl-, or methylmalonyl-lysine at position K-147. This peptide was selected because our previous proteomic datasets collected on SIRT5-KO and WT mouse brains identified this specific lysine residue in ADT1 as modified by succinylation and malonylation ^19^. We analyzed a mixture of synthetic light and heavy stable isotope-labelled peptides with malonylated, succinylated, and methylmalonylated lysine using targeted assays with ion mobility (prm-PASEF) on a Bruker timsTOF HT system. This enabled us to explore the reverse phase-liquid chromatography, ion mobility, and MS/MS behaviors of these modified peptides. The total ion chromatogram revealed three distinct peaks, with retention times at 8.45 min, 8.88 min, and 9.70 min (Fig. 2A). Matching to single peptide analysis revealed that the malonylated, succinylated, and then methylmalonylated peptides eluted in that order (Supplementary Fig. 1). Then, four ions were resolved on the ion mobility dimension with 1/K_0_ at 0.915 Vs/cm^2^ and 0.920 Vs/cm^2^ (Fig. 2B). The pair with 1/K_0_ = 0.915 Vs/cm^2^ were the precursor ions at m/z 637.82 and m/z 642.82, corresponding to the light and heavy malonylated peptides forms, respectively. The ions arriving at 1/K_0_ = 0.920 Vs/cm^2^ were the precursor ions at m/z 644.83 and m/z 649.83, that are the light and heavy forms, respectively, of the isobaric succinylated and methylmalonylated peptides.

**Figure 2.**
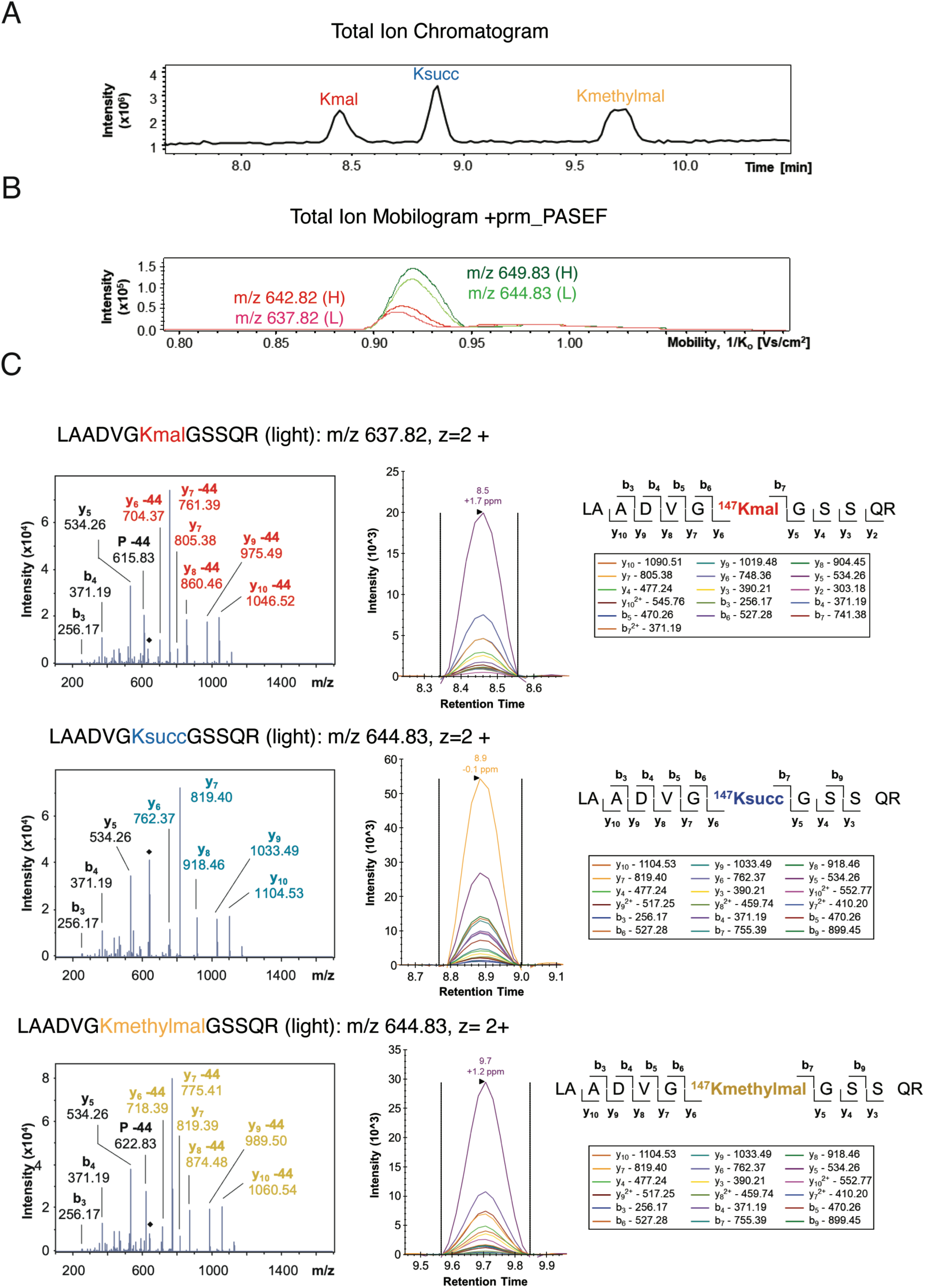
Confident differentiation of synthetic acylated peptides by LC-MS/MS analysis. A mixture of synthetic malonylated, succinylated, and methylmalonylated LAADVG^147^**Kacyl**GSSQR peptides from mouse adenine nucleotide translocator 1 (ADT1; UniProt ID: P48962), including both light and heavy stable isotope-labelled forms, was analyzed on an Evosep One liquid chromatography system coupled to a timsTOF HT mass spectrometer (Bruker Daltonics) operated in prm-PASEF mode. (A) Total ion chromatogram. Malonylated, succinylated, and methylmalonylated peptides display distinct retention times. (B) Total ion mobilogram. Trapped ion mobility spectrometry (TIMS) enabled the separation of the malonylated precursor ions at m/z 637.82 (z = 2; light form, L) and m/z 642.85 (z = 2; heavy form, H) from the isobaric succinylated and methylmalonylated precursor ions at m/z 644.83 (z = 2; L) and m/z 649.83 (z = 2; H). (C) MS/MS spectra and extracted ion chromatograms (XICs) of the light acylated precursor ions at m/z 637.82 (z = 2) corresponding to the malonylated form and at m/z 642.85 (z = 2) corresponding to the isomeric succinylated and methylmalonylated forms. The malonylated, succinylated, and methylmalonylated peptide forms were analyzed individually. On the MS/MS spectra, fragment ions displayed in color contain the PTM group and enable confident identification and site localization. Neutral loss (−44 Da corresponding to CO_2_) is observed for the malonylated and methylmalonylated species.

To investigate the fragmentation pattern of the different acylated forms, targeted analysis in prm-PASEF mode were conducted on each individual peptide, targeting malonylated precursor ion at m/z 637.82 (z = 2), succinylated and methylmalonylated precursor ion at m/z 644.83 (z = 2). The MS/MS spectra and extracted ion chromatograms (XICs) of the modified peptides are displayed on Fig. 2C. One notable observation was the detection of a neutral loss of 44 Da (CO_2_) for the fragment ions containing the malonyl and methylmalonyl group, e.g. y_6_ to y_10_ ions. Similar fragmentation pattern was observed: intact y_7_ ion at m/z 819.40 was the most intense ion for the succinylated precursor ion, and y_7_ -44 ion at m/z 761.39 and m/z 775.41 for the malonylated and methylmalonylated precursor ions, respectively. Intact y_7_ ion at m/z 805.38 and m/z 819.39 was also detected for the two latest forms. The second most intense fragment ion was y_5_ ion at m/z 534.26, that does not contain the PTM group. The MS/MS spectra and XICs showed that near complete y- and b-ion series were detected, including ions containing the modification, enabling confident PTM site localization.

Together, this demonstrates that methylmalonyl-, malonyl-, and succinyl-modified peptides can be differentiated and identified with high confidence using liquid chromatography-tandem mass spectrometry (LC-MS/MS) assays.

### Methylmalonyl proteome mapping in mouse brain reveals neuronal and myelin-associated proteins

Leveraging the cross-reactivity of the anti-malonyl-lysine antibodies and the capabilities of LC-MS/MS to differentiate the different acylated peptides, we investigated the brain methylmalonylome and its dynamic remodeling upon SIRT5 deletion using previously collected proteomic data (Fig. 3A) ^19^. Briefly, whole brain tissues were collected from 18-month-old SIRT5-KO and WT mice, proteins were digested with trypsin, and proteolytic peptides were subjected to parallel malonylated or succinylated peptide enrichments before nanoLC-MS/MS analysis using label-free data-independent acquisition (DIA) on a Thermo Orbitrap Eclipse Tribrid platform. Finally, DIA data were processed using the library-free directDIA software. Here, the DIA dataset collected on malonylated peptide enrichments were re-processed using directDIA, now defining lysine methylmalonylation as variable modification. Using this approach, we identified 44 high-confidence methylmalonylated peptides, corresponding to 43 unique PTM sites, across 41 proteins (Supplementary Table S1). One identified methylmalonylated peptide was LAAD^147^**Kmethylmal**GSSQR from ADT1. As we previously reported the identification of the malonylated LAAD^147^**Kmal**GSSQR and succinylated LAAD^147^**Ksucc**GSSQR in the mouse brain specimens, this allowed us to assess their chromatographic and fragmentation behavior relative to the synthetic reference peptides (Fig. 2 and 3B). Despite being acquired on independent LC-MS/MS platforms operating in different modes, the endogenous and heavy-labelled modified peptides demonstrated equivalent fragmentation ion intensity profiles, with y_5_ ion being the most intense ion for the malonylated and methylmalonylated precursor ions and y_7_ ion for the succinylated one. Concordant elution order was also observed as the malonylated peptide eluted first at 47.7 min, then the succinylated peptide at 49.4 min, and the methylmalonylated peptide eluted the latest at 52.1 min. This supports unambiguous identification of methylmalonylated peptides using this strategy.

**Figure 3.**
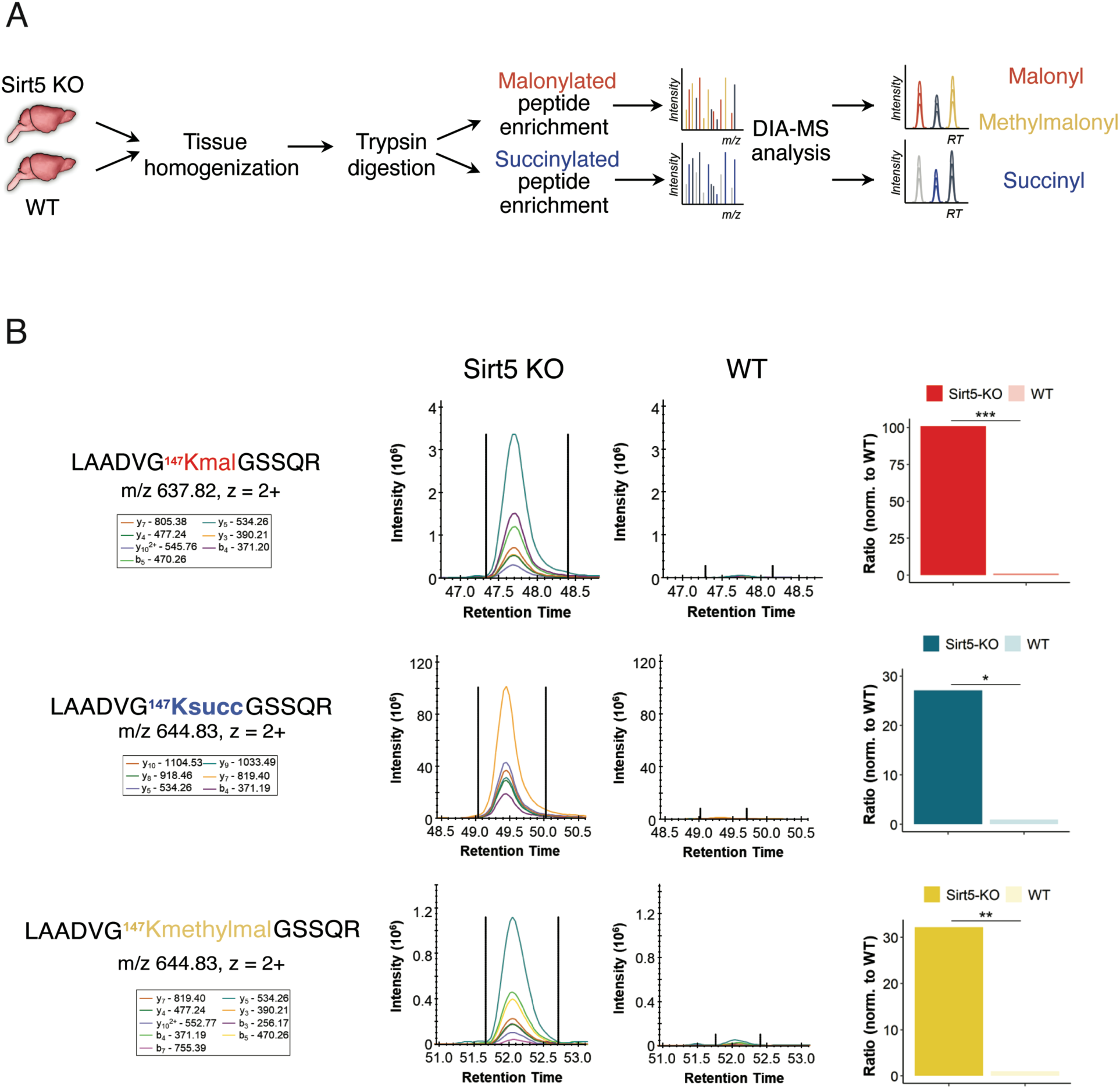
Proteome-wide mapping of lysine methylmalonylation in mouse brain. (A) Experimental workflow for methylmalonyl proteomics. We previously reported the malonylome and succinylome of wild-type (WT) and SIRT5 knockout (KO) mice (n = 4 each) ^21^; datasets available as MassIVE ID MSV000089606 and ProteomeXchange ID: PXD034275). Briefly, brain tissues from wild-type (WT) and Sirt5 knockout (KO) mice were homogenized, digested with trypsin, and subjected to modified peptide enrichment using PTMScan beads coated with antibodies recognizing lysine malonylation or lysine succinylation (Cell Signaling Technologies). Enriched peptides were analyzed by data-independent acquisition (DIA-MS) on an Orbitrap Eclipse system (Thermo Fisher Scientific) and data were processed using directDIA, which relies on a library-free strategy, directly identifying and quantifying modified peptides from the DIA data. To map the lysine methylmalonylome, the DIA-MS dataset collected on malonylated peptide enrichment was re-processed with directDIA, now searching for lysine methylmalonylation. (B) Representative extracted ion chromatograms of LAADVG^147^**Kacyl**GSSQR (malonylated precursor ion at m/z 637.82, z = 2; succinylated and methylmalonylated precursor ion at m/z 644.83, z = 2) of the mouse adenine nucleotide translocator 1 (ADT1) with a modification on Lys-147 in a SIRT5-KO and WT replicate. Statistical analysis confirmed the significant increased malonylation, succinylation and methylmalonylation level of this peptide in SIRT5-KO brain compared with WT controls (*** p-value = 4.2e-9 for malonylation, * q-value = 2.3e-2 for succinylation, ** p-value = 5.5e-5 for methylmalonylation).

Comparison of SIRT5-KO and WT brains revealed six significantly altered methylmalonylated peptides (p-value < 0.05 and log_2_ (SIRT5-KO vs WT) > 0.58), with increased methylmalonylation level at five unique sites, supporting a role for SIRT5 in constraining this modification *in vivo*. The most significant change was Lys-147 of the ADP/ATP translocase 1 (ADT1), that presented a 32-fold enrichment (p-value at 5.5e-5 and q-value at 0.01) upon SIRT5 deletion (Fig. 3B). Interestingly, we previously reported that Sirt5 also targets this site when malonylated (101-fold enrichment) and succinylated (27-fold enrichment). These results provide the first proteome-wide map of methylmalonylation in the mammalian brain and establish its susceptibility to Sirtuin 5 regulation.

Next, to assess whether methylmalonylation displays distinct biological associations relative to its isobaric counterpart succinylation, we compared gene ontology (GO) enrichment profiles between the two modifications. This analysis revealed that proteins related to cellular respiration, presenting an oxidoreductase activity, and located to mitochondria were subjected to both lysine methylmalonylation and succinylation. However, despite their chemical similarity, differences in enriched biological processes and protein classes were observed between the two modifications. Methylmalonylated proteins showed enrichment across several GO classifications, including biological processes related to purine metabolism, molecular functions such as angiostatin binding, and cellular compartments associated with the myelin sheath (Fig. 4A–C). Strikingly, several structural components of myelin and axons, such as Neurofilament medium polypeptide (NEFM), Neurofilament light polypeptide (NEFL), and Myelin Basic Protein (MBP), were detected as methylmalonylated in mouse brain (Fig. 4C). These distinct enrichment patterns suggest that methylmalonylation and succinylation occupy partially distinct proteomic landscapes and further support the ability of our workflow to discriminate between these isobaric modifications.

**Figure 4.**
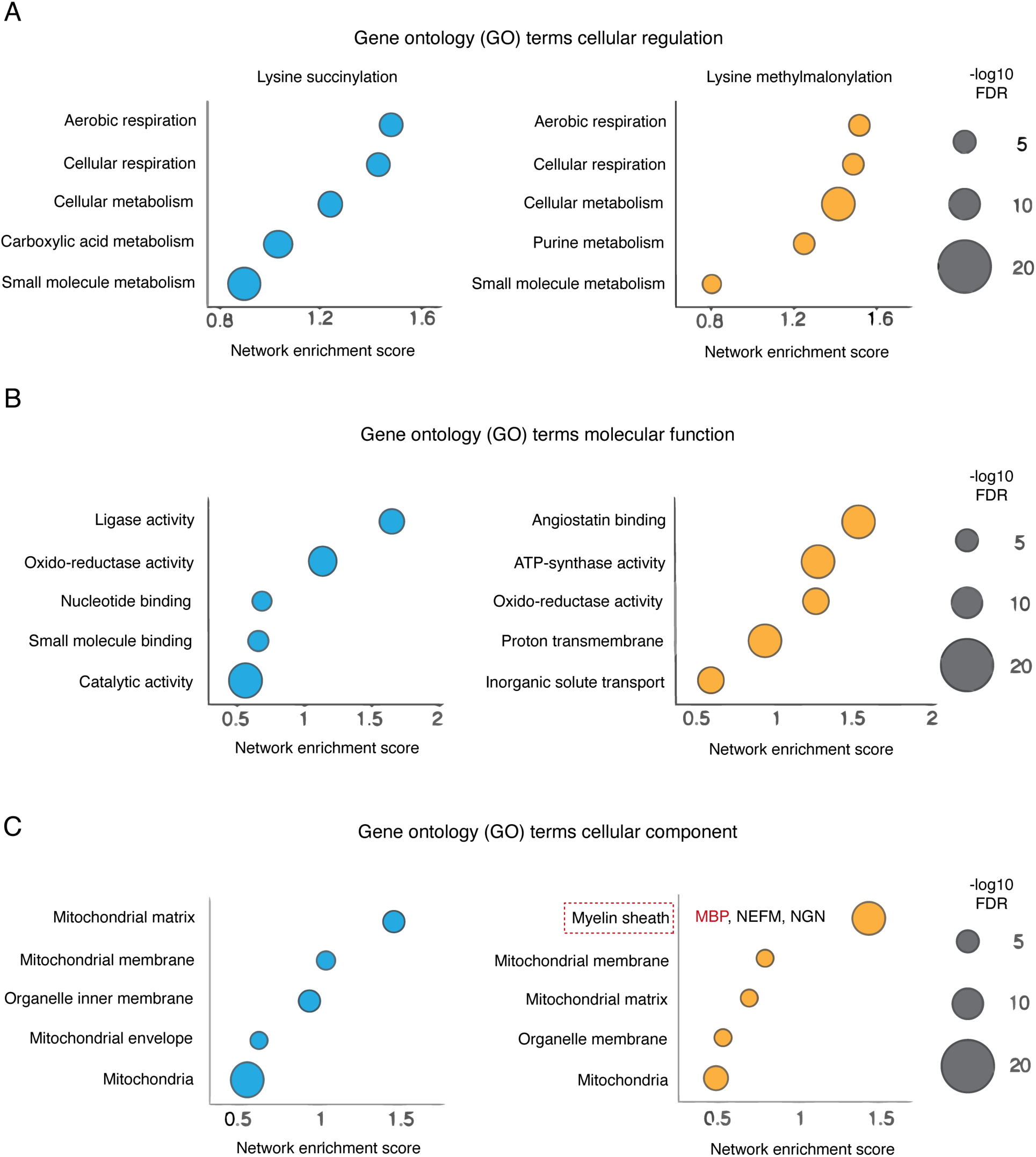
Comparative gene ontology analysis distinguishes methylmalonylated from succinylated proteome. (A–C) Comparative gene ontology (GO) enrichment analysis of proteins modified by methylmalonylation (right, in yellow) versus succinylation (left, in blue). Bubble plots illustrate enriched biological processes, molecular functions, and cellular compartments for each modified sub-proteome.

### Methylmalonylation targets myelin basic protein and disrupts its lipid-binding capacity

Given the central role of MBP in myelin membrane compaction, we examined the distribution of the methylmalonylation sites across the protein sequence (Fig. 5A). One site was Lys-129 on isoform 4 (21.5-kDa) and isoform 6 (17-kDa-a) or Lys-103 on isoform 5 (18.5-kDa) and isoform 8 (14-kDa), which was further identified as a SIRT5 target in mouse brain, and the second site was Lys-83 on isoform 4 (21.5-kDa), isoform 6 (17-kDa-a), isoform 9 and isoform 11 (19.7-kDa). Importantly, none of these residues were detected as succinylated from the succinylated peptide enrichment analysis, supporting novel methylmalonylation site identification. Methylmalonylated Lys-83 is located within the lipid-binding region responsible for mediating MBP–membrane interactions. These findings raised the possibility that negatively charged methylmalonyl modifications could directly influence MBP ability to associate with myelin membranes.

**Figure 5.**
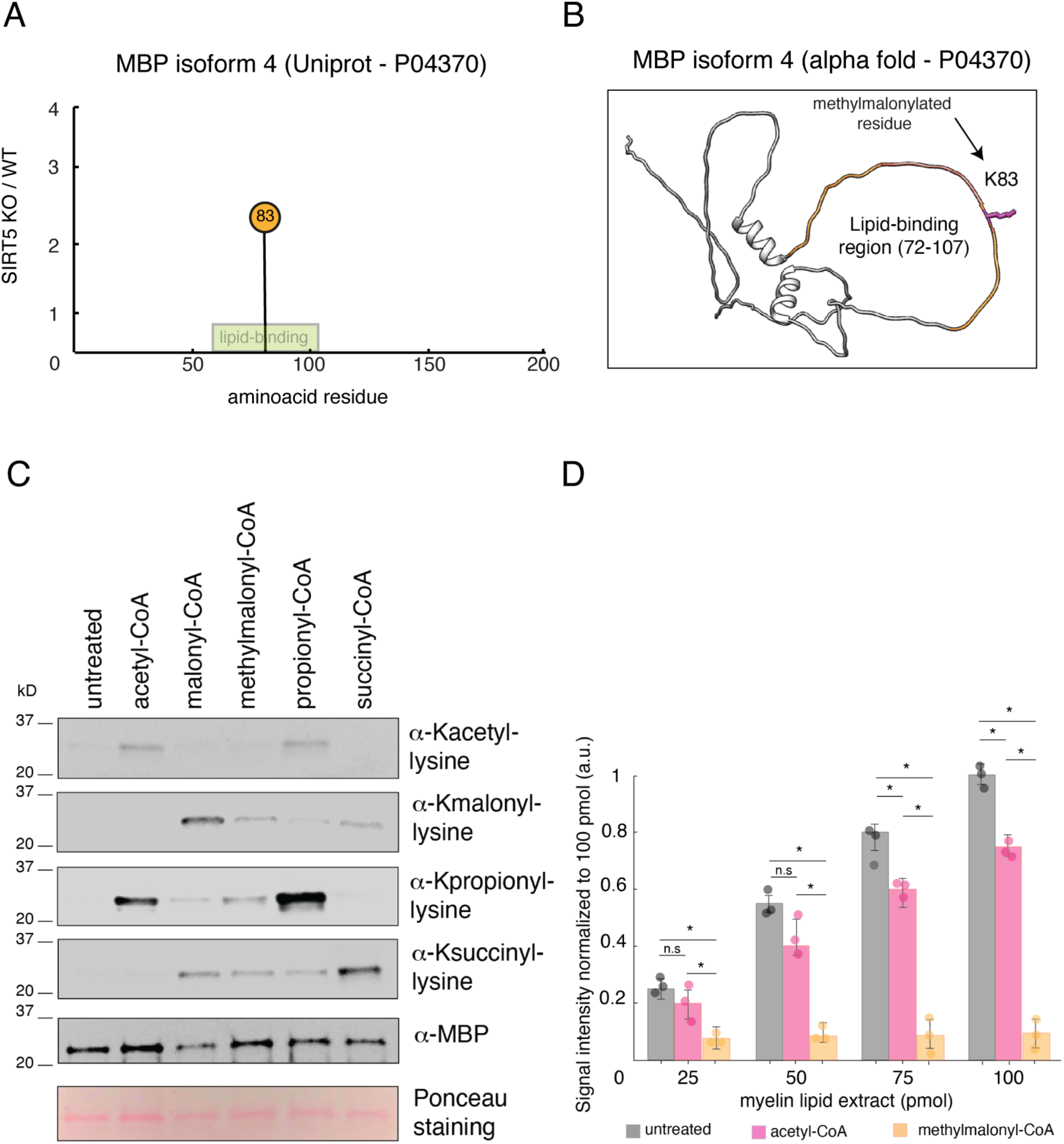
Methylmalonylation impairs lipid binding of myelin basic protein. (A) Lollipop plot showing the distribution of methylmalonylation sites along the sequence of the isoform 4 of MBP. A methylmalonylated lysine (K83) was identified (present on several MBP isoforms) within the lipid-binding region (residues 72–107) that mediates MBP interaction with myelin membranes. The position of the lipid-binding region is indicated. (B) Structural model of MBP highlighting the lipid-binding region (residues 72–107). The methylmalonylation site identified in the proteomic analysis on lysine 83 map within or near this region. (C) *In vitro* acylation of recombinant MBP using acetyl-CoA, malonyl-CoA, methylmalonyl-CoA, propionyl-CoA, or succinyl-CoA. Immunoblot analysis confirms incorporation of the corresponding lysine acyl modifications. (D) Quantification of MBP binding to myelin lipid extracts in myelin lipid overlay assays. Binding signal intensity was quantified and normalized to untreated MBP. Data are presented as mean ± SEM from three independent replicates, with individual data points shown. Statistical p values indicated in the figure (*p < 0.05).

Because MBP association with myelin membranes relies on electrostatic interactions between positively charged lysine residues and negatively charged membrane lipids, the addition of negatively charged acyl groups such as methylmalonylation could destabilize these interactions. Notably, one methylmalonylated residue identified in our proteomic analysis map was found within the lipid-binding region of MBP (residues 72–107), suggesting a potential mechanism by which this modification could impair MBP–membrane association (Fig. 5A).

To test this hypothesis, recombinant MBP was modified *in vitro* using acetyl-CoA, malonyl-CoA, methylmalonyl-CoA, or succinyl-CoA to generate defined acylated forms of the protein. Successful modification was confirmed by immunoblotting with antibodies recognizing the corresponding lysine acylations (Fig. 5C). We next assessed the ability of these modified proteins to bind myelin lipids using lipid overlay assays. Unmodified MBP displayed robust dose-dependent lipid binding. MBP acetylation, which neutralizes lysine charge without introducing a negative charge, produced a modest ∼1.3–1.5-fold reduction in lipid binding compared to untreated MBP. In contrast, the introduction of negatively charged acyl modifications—particularly methylmalonylation—consistently impaired MBP binding to myelin lipids, resulting in an approximate ∼3–6-fold decrease in binding relative to the untreated protein across the range of myelin lipid extract concentrations tested (Fig. 5D).

## Discussion

Lysine methylmalonylation has long been suspected to occur in mammalian tissues exposed to carbon stress, but its study has been hindered by analytical obstacles ^5,7,9^. Unlike other lysine acylations, no commercially available reagents exist that selectively enrich methylmalonyl-lysine, and its mass shift (+100.016 Da) is indistinguishable from succinyl-lysine by conventional MS/MS ^19^. As a result, methylmalonylation has been largely undetectable in standard acyl-proteomics workflows. Here, we present a robust strategy for the detection and confident identification of methylmalonylated peptides in complex proteomes, overcoming an analytical barrier caused by their low abundance and isobaricity with succinyl-lysine. By leveraging the cross-reactivity of anti-malonyl antibodies, high-sensitivity proteomic assays on modern high-resolution accurate-mass instruments, and efficient library-free algorithm for DIA data analysis, this workflow enabled us to generate the first proteome-wide map of lysine methylmalonylation in the mammalian brain. Our biochemical validation confirmed that anti-malonyl antibodies recognize methylmalonylated lysines but not succinylated counterparts, providing a practical basis for enrichment despite the lack of modification-specific reagents (Fig. 1). Synthetic ADT1 peptides carrying malonyl-, succinyl-, or methylmalonyl-lysine demonstrated reproducible analytical features, including retention time, ion mobility, and fragmentation profiles, differentiating these three acylations and supporting confident PTM identification and site localization (Fig. 2). Together, these benchmarks provided strong orthogonal evidence that site-specific identification of lysine methylmalonylation can be achieved with confidence.

Applying this workflow to WT and SIRT5-KO mouse brain tissues revealed 44 identified methylmalonylated peptides across 41 proteins (Supplementary Table S1). Gene ontology analysis indicated strong enrichment in neuronal structural and myelin-associated proteins, including MBP, NEFM, and NEFL, as well as mitochondrial proteins such as ADT1. Several sites were significantly increased in SIRT5 knockout brains, consistent with the established desuccinylase and demalonylase activities of this mitochondrial deacylase. These results place methylmalonylation at the intersection of neuronal structure and energy metabolism, raising the possibility that it contributes to both myelin stability and mitochondrial function under metabolic stress (Fig. 4).

Among all identified proteins, MBP was particularly notable because of its role in compact myelin formation. One methylmalonylation site clustered within positively charged domains required for lipid binding (Fig. 5A). Functional assays confirmed that *in vitro* methylmalonylation of MBP impairs its ability to interact with myelin lipids, mirroring the disruptive effects of malonylation and succinylation (Fig. 5B & C). These findings provide a mechanistic link between methylmalonylation and destabilization of myelin structure, with potential implications for both inborn errors of metabolism and age-associated demyelinating disorders.

More broadly, this work highlights a generalizable strategy for resolving isobaric acyl modifications that lack specific enrichment tools. Similar analytical blind spots exist for other acylations; for example, di-acetyl- and propionyl-lysine are also isobaric (+42 Da) and are co-enriched by pan-acyl antibodies, which likely obscures the true prevalence of propionylation ^24^. Our approach provides a template for addressing such challenges and uncovering previously unrecognized PTMs.

In conclusion, we present a novel and straightforward workflow that enables confident detection, quantification, and site localization of lysine methylmalonylation in complex tissues. This framework reveals methylmalonylation as a widespread modification in the brain, regulated by SIRT5, and capable of impairing MBP function. Future work should determine whether methylmalonylation accumulates under physiological carbon stress or in disease states such as methylmalonic acidemia, neurodegeneration, or aging. With the development of methylmalonyl-specific reagents, the strategy described here will accelerate systematic exploration of this modification and its impact on cellular metabolism and structural integrity.

## Acknowledgements

This work was supported by the NIH/NCRR shared instrumentation grant S10 OD028654 (B.S.) and by the Glenn Foundation for Medical Research (E.V.).

## Author contributions

J.L.-G., J.B., and G.V.-H. contributed equally to this work. *Conceptualization:* J.L.-G. and J.B. envisioned the enrichment of methylmalonylated peptides using anti-malonyl antibodies, in conjunction with B.S. and E.V. *Methodology*: J.B. and B.S. designed the analytical strategy to discriminate isobaric peptides. J.L.-G. and G.V.-H., with input from E.V., designed the in vitro MBP lipid-binding assays. Investigation: J.L.-G., J.B., and R.R. performed sample preparation, peptide enrichment, and proteomic analyses. J.B., R.Z., and B.S. conducted LC–MS/MS experiments. J.B. performed ion mobility experiments with design input from B.S. J.L.-G. and G.V.-H. performed recombinant protein acylation and MBP lipid-binding assays with input from E.V. *Formal Analysis*: J.L.-G. and J.B. performed primary data analysis. All authors contributed to data interpretation and discussion. *Writing* J.L.-G. and J.B. wrote the first draft with help of all authors *– Review & Editing:* All authors. Supervision: B.S., E.V. Funding Acquisition: B.S., E.V.

## Conflict of Interest

B.S. serves on the Proteomics Advisory Board of MOBILion Systems, Inc. The other authors declare that they have no known competing financial interests or personal relationships that could have appeared to influence the work reported in this paper.

## Abbreviations

ACN: acetonitrile
ADT1: adenine nucleotide translocator 1 (Slc25a4)
BCAA: branched-chain amino acid
DIA: data-independent acquisition
FA: formic acid
GO: Gene Ontology
IEM: inborn error of metabolism
LC–MS/MS: liquid chromatography-tandem mass spectrometry
MBP: myelin basic protein
MMA: methylmalonic acidemia
ncKac: negatively charged lysine acylation
NEFL: neurofilament light polypeptide
NEFM: neurofilament medium polypeptide
prm-PASEF: parallel reaction monitoring coupled to parallel accumulation serial fragmentation
PTM: post-translational modification
SIRT5: sirtuin 5
TIMS: trapped ion mobility spectrometry
WT: wild type
XIC: extracted ion chromatogram

---

**Figure Supplementary 1.**
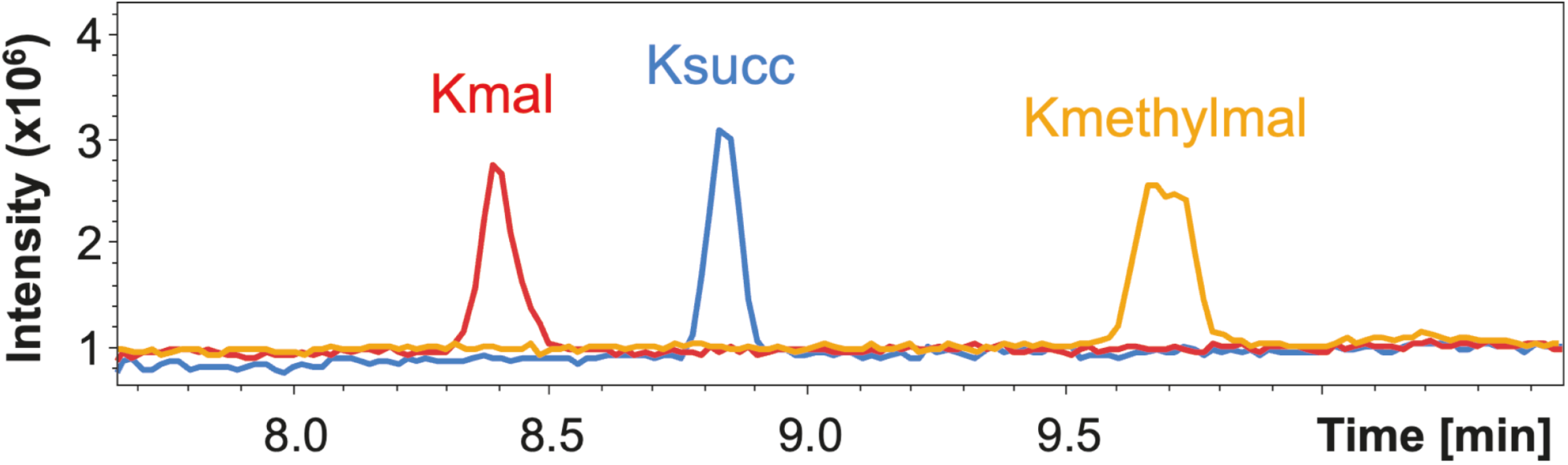
Total ion chromatograms of synthetic acylated LAADVG^147^KacylGSSQR peptide. Synthetic malonylated, succinylated, and methylmalonylated peptides LAADVG^147^KacylGSSQR from adenine nucleotide translocator 1 (ADT1; UniProt ID: P48962) were each injected individually (both light and heavy forms injected together at 10 pg each) on a timsTOF HT mass spectrometer operated in prm-PASEF mode. Peptides were eluted using a 44-min gradient at distinct retention times. Malonylated peptides eluted first at 8.40 min, then succinylated peptides at 8.83 min, and finally methylmalonylated peptides at 9.67 min.

